# Quantitative Interpretation of Transverse Spin Relaxation by Translational Diffusion in Liquids Under Arbitrary Potentials

**DOI:** 10.1101/2024.08.21.609078

**Authors:** Yusuke Okuno

## Abstract

Intermolecular spin relaxation by translational motion of spin pairs have been widely used to study properties of the biomolecules in liquids. Notably, solvent paramagnetic relaxation enhancement (sPRE) arising from paramagnetic cosolutes has gained attentions for various applications, including the structural refinement of intrinsically disordered proteins, cosolute-induced protein denaturation, and the characterization of residue-specific effective near-surface electrostatic potentials (ENS). Among these applications, the transverse sPRE rate known as Γ _2_ has been predominantly been interpreted empirically as being proportional to <*r*^-6^>_norm_. In this study, we present a rigorous theoretical interpretation of Γ _2_ that it is instead proportional to <*r*^-4^>_norm_ and provide explicit formula for calculating <*r*^-4^>_norm_ without any adjustable parameters. This interpretation is independent of the type or strength of interactions and can be broadly applied, including to the precise interpretation of ENS.

## Introduction

Solvent paramagnetic relaxation enhancement (sPRE) has been used to study protein solvent accessibility,^1-7^ to refine NMR protein structures,^8-10^ to investigate protein-cosolute interactions,^11-14^ and to characterize the electrostatic potential on small molecules and proteins.^15-26^

In many sPRE applications, the transverse sPRE rate many sPRE applications Γ_2_ is directly used to quantify electrostatics^20-25^ and for structure refinement,^8-10^ based on the Otting-LeMaster’s empirical analysis,^3, 4^ which correlates Γ _2_ with the average interspin distance <*r*^−6^>_norm_. This empirical approach has demonstrated excellent correlation with the Poisson-Boltzmann based theoretical electrostatic near-surface potentials (ENS) and the experimental ENS. However, despite its apparent success, a fully satisfactory theoretical explanation for why this approach works has yet to be established.

In previous work, we hypothesized that Γ_2_ is proportional to ⟨*r*^−4^⟩_norm_ and demonstrated its validity using a simple hard-sphere square potential model.^13^ Here, we present a comprehensive theoretical framework that establishes Γ_2_ as being proportional to ⟨*r*^−4^⟩_norm_ rather than ⟨*r*^−6^⟩_norm_. Our derivation is broadly applicable to various types of intermolecular potential, including those in confined environement, such as reverse micelle systems. While this work focuses on sPRE, the analysis is equally applicable to a wide range of intermolecular spin relaxation by translational diffusion in liquids.

### Theory

The dipole-dipole correlation function in an isotropic liquid is given by:^27, 28^

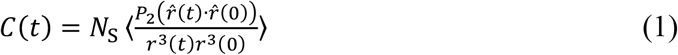

where *r* and 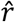 are the lengths and orientation of the interspin vector 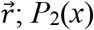 is the Legendre polynomial of degree 2; and *N*S is the number of paramagnetic cosolute molecules in the system. The number density of the cosolute *n*_S_ is related to *N*_S_ and the volume of the system *V* by 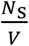.

The subscript o denotes the quantity at *t* = 0.

The spectral density *J*(*ω*) is given by the cosine transform of *C*(*t*):

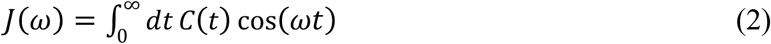

where *ω* is the angular frequency in radians.s^−1^, and is related to the spectrometer frequency (*ω*) in Hz) by *ω* = 2*πv*.

The transverse (Γ_2_) sPREs are defined as the difference in the transverse relaxation rates, respectively, of a protein nuclear spin (generally a proton) in the presence and absence of the paramagnetic cosolute.^29^ At high external magnetic field limit, Γ _2_ is related to the spectral density function *J*(*ω*) by:^30-32^

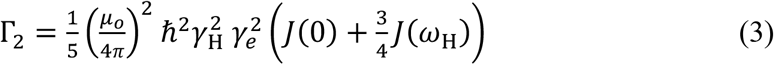

where *γ* _H_ and *γ* _e_ are the gyromagnetic ratios of the proton and electron, respectively; μ_o_ is the vacuum permittivity constant; ħ is Planck’s constant divided by 2*π*; *ω*_H_ is the angular frequency of the proton. In many instances, *J*(0) ≫ *J*(*ω*_H_) thus Eq.(3) can be approximated as

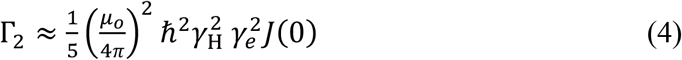

Consequently, the insights we can gain about the cosolute-protein interactions from the transverse sPRE relaxation rate is contained primarily in the details of the spectral densities *J*(0).

To relate experimentally observable sPRE relaxation rates to physical quantities, we introduced the concepts of the concentration normalized average interspin distances^11, 13, 14^

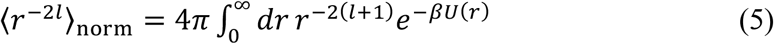

where *U*(*r*) is the potential of the mean force; *β* is the thermodynamic beta; and *l* is 2, 3, or 4. It should be emphasized that Eq.(5) is defined for any arbitrary shapes of the protein and cosolute and for arbitrary interactions between them.

If *J*(*ω*) can be measured across a wide range of the spectrometer fields, ⟨*r*^−6^⟩_norm_ can be calculated by taking the integration the spectral density function.^11^ It has also been shown that the high-frequency limit of the *ω*^*2*^*J*(*ω*) is directly related to ⟨*r*^−8^⟩_norm_.^13^

In the previous work,^13^ we hypothesized that *J*(0) is related to ⟨*r*^−4^⟩_norm_ by

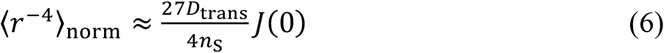

We demonstrated that Eq.(6) provide reasonable approximations for simple models involving spherically symmetric square-well and Coulombic potentials. We have also compared experimental ENS and Poisson-Boltzmann-based ENS, implemented with ⟨*r*^−4^⟩_norm_ interpretation, on ubiquitin and drkN SH3. For both systems, excellent agreements are achieved between calculated and experimental ENS similar to or slightly better than the conventionally used ⟨*r*^−4^⟩_norm_ interpretations.

Despite the apparent success of the Eq.(6), its applicability for arbitrary potentials with varying interaction strengths remained unclear. In this work, we establish a rigorous theoretical foundation of the validity of Eq.(6) and demonstrate that it applies across a wide range of systems, including those in confined environments.

In this work, we consider the simplest case where the protein and cosolutes are modeled as hardspheres, with the nuclear and electron spins located at their respective centers. The interspin vector 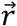 is specified by three coordinates in spherical coordinates 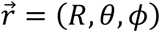. The cosolute and protein are assumed to be diffuse under the influence of the spherically symmetric potential characterized by the potential of the mean force *U*(*R*), which only depends on the center-to-center distance *R*. The cosolute is allowed to diffuse within the volume *V* specified by the contact distance *R*_C_ and the outer boundary *R*_B_ such that 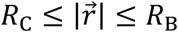. To simplify and clarify our discussion, many of the technical details of the derivations of the equations described below are provided in the Supporting Information.

Under the assumption that cosolute-protein pair is described by the Smoluchowski formalism (see Supporting Information for detail), it can be shown^13^ that the *J*(0) is given by

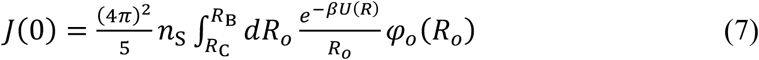

where *φ*_*o*_(*R*_*o*_) is the solution to the equation,

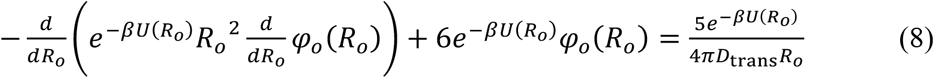

subjected to the reflective boundary conditions

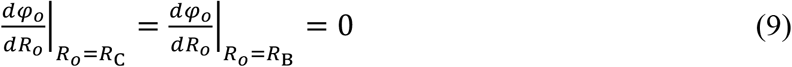

Therefore, *J*(0) can be calculated once *φ*_*o*_ is known. In practice, *φ*_*o*_ can be solved analytically only for few special cases such as force-free hardsphere (FFHS) model where *U*(*R*)=0 or when the potential of the mean force is modelled as a square-well potential.

In the case of the FFHS model, *J*(0) is given by

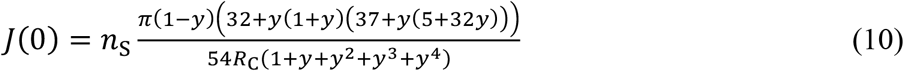

where *y* = *R*_C_/*R*_B=_. In the limit where *R*_B=_ → ∞, *J*(0) takes the simple form

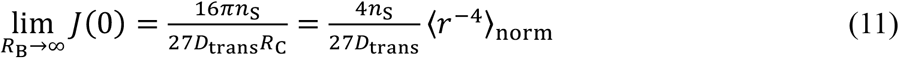

It should be pointed out that for the FFHS model with infinite volume, Eq.(11) implies that Eq.(6) is exact; however, as seen from Eq.(10), *J*(0) is not strictly equal to 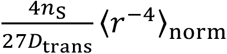 when finite volume is considered.

### Lower and Upper Bounds of *J*(0)

Since the analytical form of *φ*_*o*_ for an arbitrary potential is not known, we approximate *J*(0) by evaluating its lower and upper bounds using the complementary variational principle.^33-36^

For rigorous explanations the technique employed in this work, see refer to Ref.33, 37, 38. Let us introduce new functions *p, w, q* given by

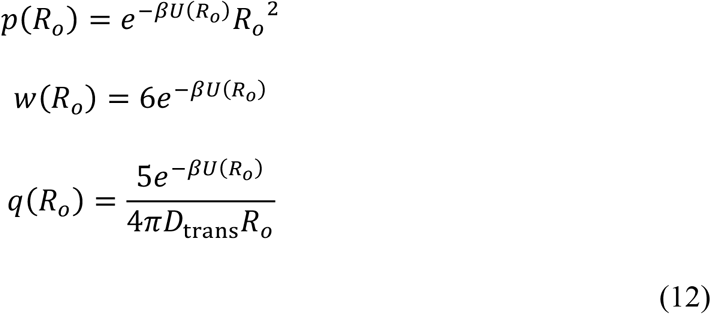

It is important to note that *p, w, q* are nonnegative functions.

Using these new functions, Eq.(8) can be written in the form of Sturm-Liouville equation

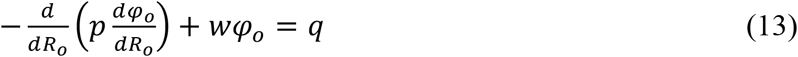

Let us introduce a functional *I*[*φ*] given by

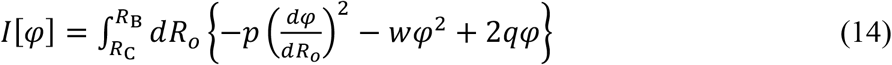

where the admissible functions of this functional are the set of all functions *φ* that have well defined first- and second derivatives and also satisfy the boundary conditions Eq.(9).

When *φ* = *φ*_*o*_,

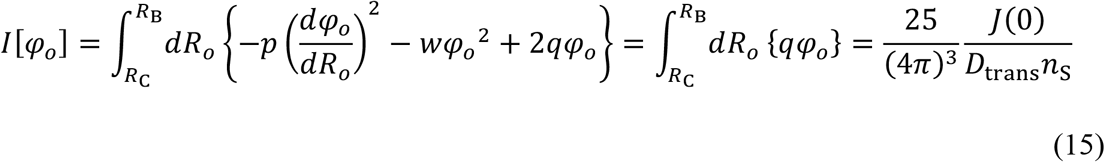

If *φ* is a function that deviates from the *φ*_*o*_ by a small amount

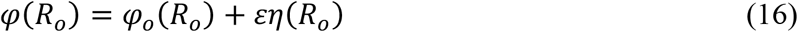

where *ε* is some small number and *η* is some arbitrary admissible function. Substituting Eq.(16) in Eq.(14), we get

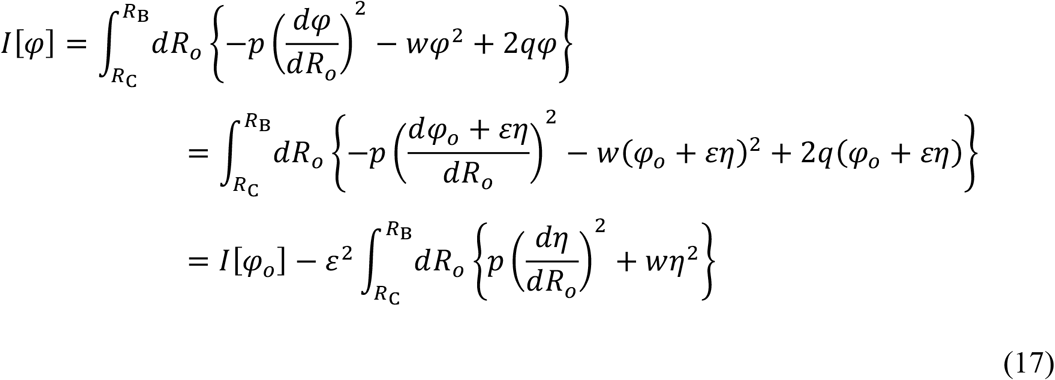

The first variation *δI* (the term linear to *ε*) is given by

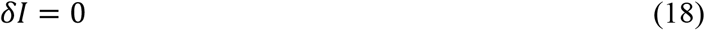

and the second variation *δ*^2^*I* (the term linear to *ε*^2^) is given by

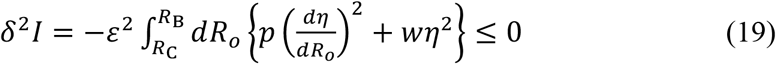

Since *I*[*φ*_*o*_] has the first variation zero and second variation negative (concave down), Eq.(18) and (19) imply that *I*[*φ*_*o*_] is the global maximum value that can attained by any admissible *φ*.^33^

To obtain the upper bound, define another function

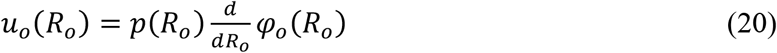

Note that *u*_*o*_ satisfies *u*_*o*_ (*R*_C_) = *u*_*o*_ (*R*_B_) = 0.

Then the Eq.(13) can be rewritten as

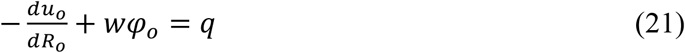

Let us introduce another functional *G*[*u*] given by

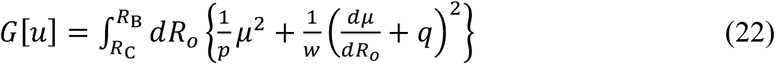

where the admissible functions is the set of all functions that have well-defined first derivative and satisfies the boundary conditions

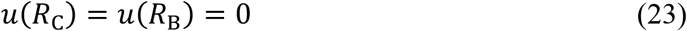

If *u* = *u*_*o*_ then we get

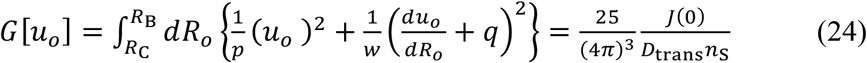

Similar to the lower bound case, we calculate the first- and second variations of *G*[*u*] by considering small deviations to *u*_*o*_. Consider the function *u* given by

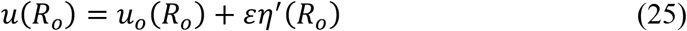

where *η*^′^ is some admissible function.

When Eq.(25) is substituted in Eq.(22), we get

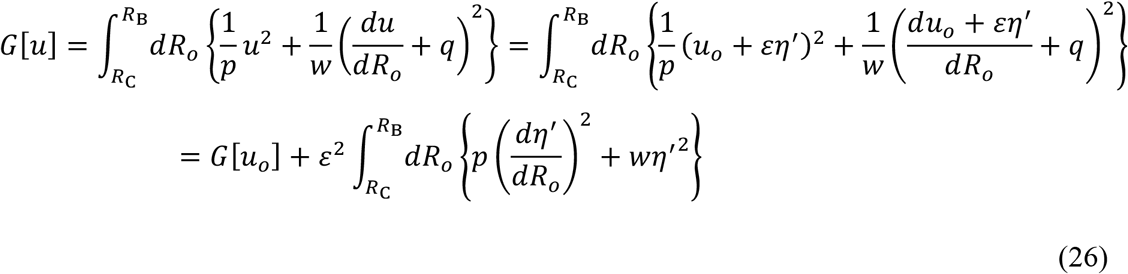

Eq.(26) implies that

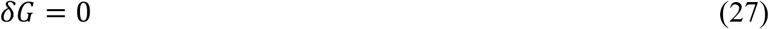

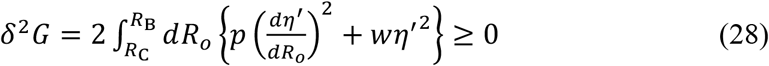

Having first variation zero and second-variation nonnegative (concave up), Eqs.(27) and (28) implies that *G*[*u*_*o*_] is the global minimum value attainable by any admissible function.^33^

To summarize our finding, we derived the following relation

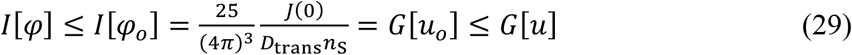

Eq.(29) provides the useful estimation of *J*(0) by choosing proper choice of the trial functions *φ* and *u*.

Here, the FFHS model (*U*(*R*)=0) was used as the trial function by setting *φ* = *φ*_FHHS_ to obtain the lower bound. Similarly, upper bound were found by substituting trivial function *u* = 0 into Eq.(22). After some rearrangement of the equation, we obtained the main finding of this work

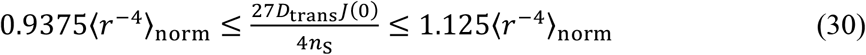

Eq.(30) suggest that 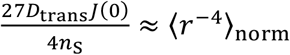 with the error up to 12.5 %. Consequently, once *D*_trans_ is acquired, a reasonable estimate of ⟨*r*^−4^⟩_norm_ can be determined without any adjustable parameters. It should be emphasized that Eq.(30) holds true for arbitrary potential of any interaction strength. Furthermore, Eq.(30) does not depend on the value of *R*_B_ (see Supporting Information and Fig.S1), making it applicable to confined environments, such as reverse-micelles, under assumption that the diffusive motion of the cosolute is described by Smoluchowski equation Eq.(S2). As a result, Eq.(30) provides the rigorous interpretation of *J*(0) for wide variety of systems. To illustrate this, we simulated the spectral densities using Lennard-Jones and Coulombic potentials and demonstrated that the inequality Eq.(30) indeed holds valid (Figure 1).

**Figure 1.**
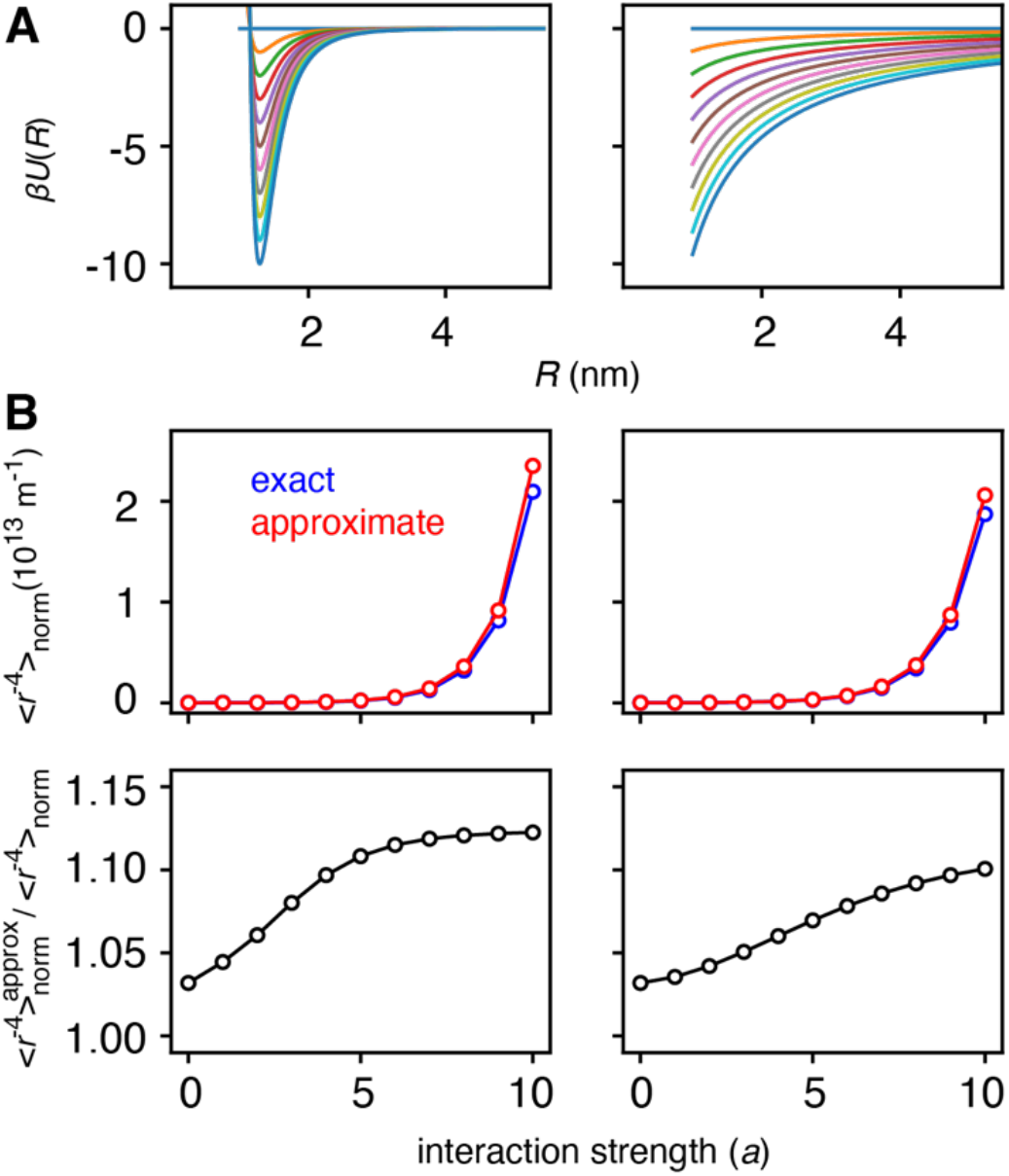
Illustration of the validity of Eq.(30). (A) The examples of the potentials used to simulate ⟨*r*^−4^⟩_norm_ and the approximate 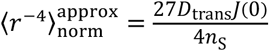. The Lennard-Jones potential of the form 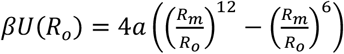 was used to generates the points on the left panel. The potential is minimum at 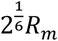. The Coulomb potential of the form 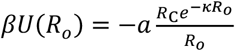 where *k* is the inverse of the Debye length was used to simulate the points on the right panel. For both potentials, *a* is the parameter used to modulate the strengths of protein-cosolute interaction. (B) The comparison between exact and approximate ⟨*r*^−4^⟩_norm_. The exact ⟨*r*^−4^⟩_norm_ were calculated from numerically evaluating the integral Eq.(5). The approximate ⟨*r*^−4^⟩_norm_ were calculated using Eq.(6). The corresponding 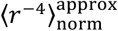 are numerically calculated by finite-difference method explained in details in the Supporting Information of the Ref. 13. The relevant parameters used for the finite-difference methods are: ∆*R* = 10^−12^ m, *M* = 120, *N* = 500 and *c* = 1.01. The values for the other parameters used in the simulations are: *R*_C_ = 1 nm, *R*_B_ = 5.8 nm, *R*_*m*_ = 1.15 nm and the Debye lengths 1/*k* = 25 nm. It should be pointed out that the numerical values of *n*_S_ and *D*_trans_ do not affect our results.

Interestingly, the upper bound found in Eq.(30) combined with the continuity of the first derivative of *φ*_*o*_ implies that 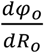 is always less than or equal to zero. That is, *φ*_*o*_ (*R*_*o*_) is a monotonically decreasing function, which may not be intuitively obvious.

## Conclusion

We have developed a new quantitative interpretation of *J*(0) for intermolecular spin relaxation, directly related to the transverse relaxation rate by considering the lower and upper bounds of *J*(0) in terms of ⟨*r*^−4^⟩_norm_. Specifically, Eq.(30) demonstrates that *J*(0) is proportional to ⟨*r*^−4^⟩_norm_, in contrast to the widely used assumption that *J*(0) is proportional to ⟨*r*^−6^⟩_norm_. Since Eq.(30) holds true for arbitrary potential, including electrostatics and van der Waals interactions, our finding provides a rigorous basis for interpreting sPRE applications on protein structure refinement and the calculation of ENS. Importantly, our approach to calculating ⟨*r*^−4^⟩_norm_ is absolute without involving any adjustable parameters. In this study, we focused on a spherically symmetric potential. Future work will investigate the validity of Eq.(30) in more complex potentials with more intricate protein and cosolute shapes.

## Acknowledgement

This work was funded by the start-up funds by Washington University in St. Louis.

