## Supporting Information for "Quantitative Interpretation of Transverse Spin Relaxation by Translational Diffusion in Liquids Under Arbitrary Potentials"

Yusuke Okuno[1]

[1]Department of Chemistry, Washington University in St. Louis, St. Louis, Missouri 63130, United States

**Derivation of Eqs.(7)-(9)**

In this Appendix, we briefly discuss the derivations of Eqs.(7)-(9). More detailed explanations can be found in Ref.1.

The time-correlation function given in Eq.(1) can be written as[1]

$$C(t) = N_{\mathrm{S}} \langle \frac{P_l(\hat{R}(t)\cdot\hat{R}(0))}{R^{l+1}(t)R^{l+1}(0)} \rangle = 4\pi n_{\mathrm{S}} \int d\vec{R} \int d\vec{R}_o \frac{Y_{l,0}(\theta,\phi)Y_{l,0}^*(\theta_o,\phi_o)}{R^{l+1}R_o^{l+1}} P(\vec{R},t|\vec{R}_o)e^{-\beta U(R)} \quad \text{(S1)}$$

where $P(\vec{R},t|\vec{R}_o)$ is the conditional probability of finding cosolute at $\vec{R}$ at $t$ given that the cosolute is at $\vec{R}_o$ at $t$=0; $Y_{l,0}(\theta,\phi)$ is the spherical harmonics $Y_{l,m}(\theta,\phi)$ with $m = 0$; $Y_{l,m}^*(\theta,\phi)$ is the complex conjugate of $Y_{l,m}(\theta,\phi)$. The integral in Eq.(S1) is taken over all possible positions and orientations of the protein-cosolute pair.

Following Hwang and Freed,[2] we approximate $P(\vec{R},t|\vec{R}_o)$ as the solution of the Smoluchowski Equation given by

$$\frac{\partial P(\vec{R},t|\vec{R}_o)}{\partial t} = D_{\mathrm{trans}} \nabla_R \cdot e^{-\beta U(R)} \nabla_R e^{\beta U(R)} P(\vec{R},t|\vec{R}_o) \quad \text{(S2)}$$

subjected to the reflective boundary conditions

$$\frac{\partial}{\partial R} e^{\beta U(R)} P(\vec{R},t|\vec{R}_o)\Big|_{R_o=R_{\mathrm{C}}} = \frac{\partial}{\partial R} e^{\beta U(R)} P(\vec{R},t|\vec{R}_o)\Big|_{R_o=R_{\mathrm{B}}} = 0 \quad \text{(S3)}$$

The gradient operator $\nabla_R$ is acting on the variable $R$ in Eq.(S2).

Corresponding adjoint Smoluchowski equation[3] is given by

$$\frac{\partial P(\vec{R},t|\vec{R}_o)}{\partial t} = D_{\mathrm{trans}} e^{\beta U(R_o)} \nabla_{R_o} \cdot e^{-\beta U(R_o)} \nabla_{R_o} P(\vec{R},t|\vec{R}_o) \quad \text{(S4)}$$

It should be emphasized that the gradient operator $\nabla_{R_o}$ is now acting on $R_o$.

Corresponding boundary conditions for adjoint Smoluchowki equation is given by

$$\frac{\partial P(\vec{R},t|\vec{R}_o)}{\partial R_o}\Bigg|_{R_o=R_{\mathrm{C}}} = \frac{\partial P(\vec{R},t|\vec{R}_o)}{\partial R_o}\Bigg|_{R_o=R_{\mathrm{B}}} = 0 \quad \text{(S5)}$$

Taking the Laplace transform of Eq.(S2), we get

$$\left(s - D_{\mathrm{trans}} e^{\beta U(R_o)} \nabla_{R_o} \cdot e^{-\beta U(R_o)} \nabla_{R_o}\right) \tilde{P}(\vec{R},s|\vec{R}_o) = -\delta(\vec{r}-\vec{r}_o) \quad \text{(S6)}$$

where $\tilde{P}(\vec{R},s|\vec{R}_o)$ is the Laplace transform of $P(\vec{R},t|\vec{R}_o)$.

The spherical symmetric form of the equation Eq.(S2) implies that the solution is of the form[4]

$$\tilde{P}(\vec{R},s|\vec{R}_o) = \frac{4\pi}{2l+1} \sum_{l=0}^{\infty} f_l\,(R,s|R_o) \sum_{m=-l}^{l} Y_{l,m}(\theta,\phi) Y_{l,m}^*(\theta_o,\phi_o) \quad \text{(S7)}$$

where $f_l(R, s|R_o)$ is a function that depends only on the distances $R$ and $R_o$.

Substituting Eq.(S6) into Eq.(S5), we obtain the equation for $f_l(R, s|R_o)$

$$\left(s - D_{\text{trans}} \frac{e^{\beta U(R_o)}}{R_o{}^2} \frac{\partial}{\partial R_o} e^{-\beta U(R_o)} R_o{}^2 \frac{\partial}{\partial R_o} - \frac{D_{\text{trans}} l(l+1)}{R_o{}^2}\right) f_l(R, s|R_o) = -\frac{2l+1}{4\pi R_o{}^2} \delta(R - R_o) \tag{S8}$$

Corresponding boundary conditions for $f_l(R, s|R_o)$ is given by

$$\left.\frac{\partial f_l(R, s|R_o)}{\partial R_o}\right|_{R_o = R_\text{C}} = \left.\frac{\partial f_l(R, s|R_o)}{\partial R_o}\right|_{R_o = R_\text{B}} = 0 \tag{S9}$$

Based on Eq.(2), we observe that

$$J(0) = 4\pi n_\text{S} \int d\vec{R} \int d\vec{R}_o \frac{Y_{l,0}(\theta,\phi) Y_{l,0}^*(\theta_o,\phi_o)}{R^{l+1} R_o^{l+1}} e^{-\beta U(R_o)} \tilde{P}\left(\vec{R}, 0\middle|\vec{R}_o\right) \tag{S10}$$

It follows from Eq.(S7) and (S10) that

$$J(0) = \frac{(4\pi)^2}{2l+1} n_\text{S} \int_{R_\text{C}}^{R_\text{B}} dR_o \int_{R_\text{C}}^{R_\text{B}} dR\, e^{-\beta U(R_o)} R^{l+1} R_o^{l+1} f_l(R, 0|R_o) \tag{S11}$$

Let us define new function $\varphi_o(R_o)$ given by

$$\varphi_o(R_o) = \int_{R_\text{C}}^{R_\text{B}} dR\, R^{l+1} f_l(R, 0|R_o) \tag{S12}$$

Substituting Eq.(S12), we obtain Eq.(7) in the main text

$$J(0) = \frac{(4\pi)^2}{2l+1} n_\text{S} \int_{R_\text{C}}^{R_\text{B}} dR_o \left\{e^{-\beta U(R_o)} R_o^{l+1} \varphi_o(R_o)\right\} \tag{S13}$$

Substituting Eq.(S12) in Eq.(S8), it is easy to see that $\varphi_o(R_o)$ satisfies the following equation corresponding to Eqs.(8)-(9) when $l = 2$,

$$D_{\text{trans}} \frac{e^{\beta U(R_o)}}{R_o{}^2} \frac{d}{dR_o} e^{-\beta U(R_o)} R_o{}^2 \frac{d}{dR_o} \varphi_o(R_o) - l(l+1)\varphi_o(R_o) = -\frac{2l+1}{4\pi} R_o{}^{-l-1} \tag{S14}$$

with the boundary conditions

$$\left.\frac{d\varphi_o}{dR_o}\right|_{R_o = R_\text{C}} = \left.\frac{d\varphi_o}{dR_o}\right|_{R_o = R_\text{B}} = 0 \tag{S15}$$

**Derivation of the first variations for $I[\varphi]$ and $G[u]$**

Let us denote some functions $p, w$, and $q$ given by

$$p(R_o) = e^{-\beta U(R_o)} R_o{}^2$$

$$w(R_o) = l(l+1) e^{-\beta U(R_o)}$$

$$q(R_o) = e^{-\beta U(R_o)} \frac{2l+1}{4\pi} R_o{}^{-l+1}$$

(S16)

These functions are exactly functions defined in Eq.(12) when $l = 2$. For our records, we keep the $l$ unspecified in this discussion, but the results in the main text is given by substituting 2 for $l$.

Using $p, w$, and $q$, Eq.(S14) can be written simply as

$$-\frac{d}{dR_o} p \frac{d\varphi_o}{dR_o} + w\varphi_o = q \tag{S17}$$

Let us introduce the functional $I[\varphi]$,

$$I[\varphi] = \int_{R_\mathrm{C}}^{R_\mathrm{B}} dR_o \left\{ -p \left( \frac{d\varphi}{dR_o} \right)^2 - w\varphi^2 + 2q\varphi \right\} \tag{S18}$$

Now we define the admissible functions of $I[\varphi]$ as any function whose first- and second-derivatives are well-defined and satisfies the boundary conditions Eq.(S15). Eq.(S18) can be also written as

$$\begin{aligned} I[\varphi] &= \int_{R_\mathrm{C}}^{R_\mathrm{B}} dR_o \left\{ -p \left( \frac{d\varphi}{dR_o} \right)^2 - w\varphi^2 + 2q\varphi \right\} \\ &= -\varphi p \frac{d\varphi}{dR_o}\bigg|_{R_o = R_\mathrm{B}} + \varphi p \frac{d\varphi}{dR_o}\bigg|_{R = R_\mathrm{C}} + \int_{R_\mathrm{C}}^{R_\mathrm{B}} dR_o \left\{ \varphi \frac{d}{dR_o} p \frac{d\varphi}{dR_o} - w\varphi^2 + 2q\varphi \right\} \\ &= \int_{R_\mathrm{C}}^{R_\mathrm{B}} dR_o \left\{ \left( \frac{d}{dR_o} p \frac{d\varphi}{dR_o} - w\varphi + q \right) \varphi + q\varphi \right\} \end{aligned} \tag{S19}$$

where integrations by parts was applied in the second line of Eq.(S19). The first two terms in Eq.(S18) vanishes for $\varphi$ by the boundary conditions Eq.(S15),

When $\varphi = \varphi_o$,

$$I[\varphi_o] = \int_{R_\mathrm{C}}^{R_\mathrm{B}} dR_o \left\{ \left( \frac{d}{dR_o} p \frac{d\varphi}{dR_o} - w\varphi_o + q \right) \varphi_o + q\varphi_o \right\} = \int_{R_\mathrm{C}}^{R_\mathrm{B}} dR_o \left\{ q\varphi_o \right\} = \frac{(2l+1)^2}{(4\pi)^3 n_\mathrm{S}} \frac{J(0)}{D_\mathrm{trans}} \tag{S20}$$

where the identity Eq.(S17) was used.

For the small variations around $\varphi_o$,

$$\varphi(R_o) = \varphi_o(R_o) + \varepsilon\eta(R_o) \tag{S21}$$

Substituting Eq.(S21) into Eq.(S19), we get

$$I[\varphi] = \int_{R_\mathrm{C}}^{R_\mathrm{B}} dR_o \left\{ -p\left(\frac{d\varphi}{dR_o}\right)^2 - w\varphi^2 + 2q\varphi \right\}$$
$$= \int_{R_\mathrm{C}}^{R_\mathrm{B}} dR_o \left\{ -p\left(\frac{d\varphi_o + \varepsilon\eta}{dR_o}\right)^2 - w(\varphi_o + \varepsilon\eta)^2 + 2q(\varphi_o + \varepsilon\eta) \right\}$$
$$= I[\varphi_o] + 2\varepsilon \int_{R_\mathrm{C}}^{R_\mathrm{B}} d\, R_o \left\{ -p\left(\frac{d\eta}{dR_o}\right)\left(\frac{d\varphi_o}{dR_o}\right) - w\varphi_o\eta + q\eta \right\}$$
$$- \varepsilon^2 \int_{R_\mathrm{C}}^{R_\mathrm{B}} dR_o \left\{ p\left(\frac{d\eta}{dR_o}\right)^2 + w\eta^2 \right\} = I[\varphi_o] - \varepsilon^2 \int_{R_\mathrm{C}}^{R_\mathrm{B}} dR_o \left\{ p\left(\frac{d\eta}{dR_o}\right)^2 + w\eta^2 \right\}$$

(S22)

The first variation $\delta I[\varphi]$ is given by

$$\frac{\delta I[\varphi]}{\varepsilon} = 2\int_{R_\mathrm{C}}^{R_\mathrm{B}} dR_o \left\{ -p\left(\frac{d\eta}{dR_o}\right)\left(\frac{d\varphi_o}{dR_o}\right) - w\varphi_o\eta + q\eta \right\}$$
$$= -2\eta p \frac{d\varphi_o}{dR_o}\bigg|_{R_o = R_\mathrm{B}} + 2\eta p \frac{d\varphi_o}{dR_o}\bigg|_{R_o = R_\mathrm{C}} + 2\int_{R_\mathrm{C}}^{R_\mathrm{B}} dR_o\, \eta \left\{ \frac{d}{dR_o} p \frac{d\varphi_o}{dR_o} - w\varphi_o + q \right\} = 0$$

(S23)

where the first two terms vanish for arbitrary $\eta$ because $\varphi_o$ satisfies the boundary conditions Eq.(S17). This shows that the first variation $\delta I[\varphi]$ is zero.

For the derivation of the upper bound, let us introduce another functional given by

$$G[u] = \int_{R_\mathrm{C}}^{R_\mathrm{B}} dR_o \left\{ \frac{1}{p}u^2 + \frac{1}{w}\left(\frac{du}{dR_o} + q\right)^2 \right\} \quad \text{(S24)}$$

When this functional is evaluated at $u = u_o$ where $u_o$ is given by

$$u_o = p\frac{d\varphi_o}{dR_o} \quad \text{(S25)}$$

following equality is obtained

$$G[u_o] = \int_{R_\mathrm{C}}^{R_\mathrm{B}} dR_o \left\{ \frac{1}{p}(u_o)^2 + \frac{1}{w}\left(\frac{du_o}{dR_o} + q\right)^2 \right\} = \int_{R_\mathrm{C}}^{R_\mathrm{B}} dR_o \left\{ p\left(\frac{d\varphi_o}{dR_o}\right)^2 + w{\varphi_o}^2 \right\}$$
$$= \int_{R_\mathrm{C}}^{R_\mathrm{B}} dR_o \left\{ \left( -\frac{d}{dR_o}\left(p\frac{d\varphi_o}{dR_o}\right) + w\varphi_o \right)\varphi_o \right\} = \int_{R_\mathrm{C}}^{R_\mathrm{B}} dR_o \{q\varphi_o\} = \frac{(2l+1)^2}{(4\pi)^3 n_\mathrm{S}} \frac{J(0)}{D_\mathrm{trans}}$$

(S26)

We define the admissible function for $G[u]$ be the continuous function with well-defined first-derivative and satisfies the boundary conditions

$$u(R_\mathrm{C}) = u(R_\mathrm{B}) = 0 \quad \text{(S27)}$$

When we consider the small variations around $u_o$,

$$u(R_o) = u_o(R_o) + \varepsilon\eta'(R_o) \tag{S28}$$

we get

$$\begin{aligned} G[u] &= \int_{R_\mathrm{C}}^{R_\mathrm{B}} dR_o \left\{\frac{1}{p}u^2 + \frac{1}{w}\left(\frac{du}{dR_o} + q\right)^2\right\} = \int_{R_\mathrm{C}}^{R_\mathrm{B}} dR_o \left\{\frac{1}{p}(u_o + \varepsilon\eta')^2 + \frac{1}{w}\left(\frac{du_o + \varepsilon\eta'}{dR_o} + q\right)^2\right\} \\ &= G[u_o] + 2\varepsilon \int_{R_\mathrm{C}}^{R_\mathrm{B}} dR_o \left\{\frac{1}{p}u_o\eta' + \frac{1}{w}\left(\frac{du_o}{dR_o} + q\right)\frac{d\eta'}{dR_o}\right\} + \varepsilon^2 \int_{R_\mathrm{C}}^{R_\mathrm{B}} dR_o \left\{\frac{1}{p}{\eta'}^2 + \frac{1}{w}\left(\frac{d\eta'}{dR_o}\right)^2\right\} \\ &= G[u_o] + \varepsilon^2 \int_{R_\mathrm{C}}^{R_\mathrm{B}} dR_o \left\{p\left(\frac{\partial\eta'}{\partial R_o}\right)^2 + w{\eta'}^2\right\} \end{aligned} \tag{S29}$$

The first variation $\delta G[u]$ is explicitly evaluated as

$$\begin{aligned} \delta G[u]/\varepsilon &= 2\int_{R_\mathrm{C}}^{R_\mathrm{B}} dR_o \left\{\frac{1}{p}u_o\eta' + \frac{1}{w}\left(\frac{du_o}{dR_o} + q\right)\frac{d\eta'}{dR_o}\right\} = 2\int_{R_\mathrm{C}}^{R_\mathrm{B}} dR_o \left\{\frac{d\varphi_o}{dR_o}\eta' + \varphi_o\frac{d\eta'}{dR_o}\right\} \\ &= 2\varphi_o\eta'|_{R_o=R_\mathrm{B}} - 2\varphi_o\eta'|_{R=R_\mathrm{C}} + 2\int_{R_\mathrm{C}}^{R_\mathrm{B}} dR_o\,\eta' \left\{(-\varphi_o + \varphi_o)\frac{d\eta'}{dR_o}\right\} = 0 \end{aligned} \tag{S30}$$

Since $\eta'$ also satisfies the boundary conditions Eq.(S27), the first two terms vanish.

**Analytic expression of $\varphi_\mathrm{HS}$**

When $U(R_o) = 0$, the Eq.(S14) is simplified to

$$-\frac{d}{dR_o}{R_o}^2\frac{d}{dR_o}\varphi_\mathrm{HS} + l(l+1)\varphi_\mathrm{HS} = \frac{2l+1}{4\pi D_\mathrm{trans}}{R_o}^{-l+1} \tag{S31}$$

We found the solution to Eq.(S31) as

$$\varphi_\mathrm{HS}(R_o) = s_l(R_o) + \xi_l(R_o) \tag{S32}$$

where

$$D_\mathrm{trans}R_\mathrm{B}^{l-1}s_l(R_o) = \frac{1}{8\pi}\frac{2l+1}{2l-1}(x - xy + y)^{-l+1} \tag{S34}$$

$$\begin{aligned} &D_\mathrm{trans}R_\mathrm{B}^{l-1}\xi_l(R_o) \\ &\quad= \frac{1}{8\pi}\frac{(2l+1)(l-1)}{l(l+1)(2l-1)}\frac{(x - xy + y)^{-l}}{1 - y^{2l+1}}\Big((l+1)(1-y^2)(x - xy + y)^{2l+1} \\ &\quad- ly^2\left(1 - y^{2l-1}\right)\Big) \end{aligned} \tag{S35}$$

where $x(R_o) = \frac{R_o - R_C}{R_B - R_C}$ and $y = R_C/R_B$.

Substituting $\varphi_{HS}$ as a trial function of $I[\varphi]$,

$$I[\varphi_{HS}] = \int_{R_C}^{R_B} dR_o \left\{ -p\left(\frac{d\varphi}{dR_o}\right)^2 - w\varphi^2 + 2q\varphi \right\}$$
$$= \int_{R_C}^{R_B} dR_o\, e^{-\beta U(R_o)} \left\{ -R_o{}^2 \left(\frac{ds_l}{dR_o}\right)^2 - l(l+1)s_l^2 + \frac{(2l+1)}{2\pi D_{\mathrm{trans}}} R_o{}^{-l+1} s_l \right\}$$
$$+ \int_{R_C}^{R_B} dR_o\, e^{-\beta U(R_o)} v_l(R_o)$$

(S36)

where

$$v_l(R_o) = \left\{ -R_o{}^2 \left(\frac{d\xi_l}{dR_o}\right)^2 - 2R_o{}^2 \frac{d\xi_l}{dR_o}\frac{ds_l}{dR_o} - l(l+1)\{\xi_l^2 + 2\xi_l s_l\} + \frac{(2l+1)}{2\pi D_{\mathrm{trans}}} R_o{}^{-l+1} \xi_l \right\} \quad \text{(S37)}$$

It can be shown that $v_l(R_o)$ is always nonnegative for $l = 2$ (see Fig.S1) and the first term in Eq.(S36) can be written as

$$-\left(\frac{1}{8\pi}\right)^2 \frac{(2l+1)^2}{(2l-1)^2 (D_{\mathrm{trans}})^2} (2l^2 - 9l + 5) \int_{R_C}^{R_B} dR_o\, e^{-\beta U(R_o)} R_o{}^{-2l+2}$$
$$= \frac{2}{(8\pi)^3} \frac{(2l+1)^2}{(2l-1)^2 (D_{\mathrm{trans}})^2} (2l^2 - 9l + 5) \langle R^{-2l} \rangle_{\mathrm{norm}}$$

(S38)

It follows that

$$\frac{2}{(8\pi)^3} \frac{(2l+1)^2}{(2l-1)^2} \{-(2l^2 - 9l + 5)\langle R^{-2l} \rangle_{\mathrm{norm}}\} \leq I[\varphi_{HS}]$$

(S39)

We now find the upper bound by considering the simplest admissible function that trivially satisfies the boundary condition Eq.(S27)

$$\mu_{\mathrm{trial}} = 0 \quad \text{(S40)}$$

Substituting Eq.(S40) into Eq.(S24), we get

$$G[\mu_{\mathrm{trial}}] = \int_{R_C}^{R_B} dR_o \left\{ \frac{1}{w} q^2 \right\} = \left(\frac{1}{4\pi}\right)^2 \frac{(2l+1)^2}{l(l+1)} \int_{R_C}^{R_B} dR_o \left\{ e^{-\beta U(R_o)} R_o^{-2l+2} \right\}$$
$$= \frac{1}{(4\pi)^3} \frac{(2l+1)^2}{l(l+1)(D_{\mathrm{trans}})^2} \langle R^{-2l} \rangle_{\mathrm{norm}}$$

(S41)

In summary, we get the inequality

$$\frac{-(2l^2-9l+5)}{4(2l-1)^2(D_{\text{trans}})^2}\langle R^{-2l}\rangle_{\text{norm}} \le \frac{J(0)}{D_{\text{trans}}n_S} \le \frac{1}{l(l+1)(D_{\text{trans}})^2}\langle R^{-2l}\rangle_{\text{norm}} \tag{S42}$$

In particular, we get Eq.(30) for $l = 2$.

**Monotonically decreasing properties of $\varphi_o(R_o)$**

First, it should be pointed out that our derivation of the upper-bound Eqs.(S24)-(S27), (S40)-(S41) holds for any intervals $(x_n, x_{n+1})$ that satisfies the boundary conditions

$$\left.\frac{d\varphi_o}{dR_o}\right|_{R_o=x_n} = 0 \tag{S43}$$

Since $\frac{d\varphi_o}{dR_o}$ is continuous $\varphi_o(R_o)$ being the solution to the second-order differential equation, any change in sign of $\frac{d\varphi_o}{dR_o}$ must pass the point where $\frac{d\varphi_o}{dR_o} = 0$. Let such points be $x_1 < x_2 < \cdots$. That is, the sign of $\frac{d\varphi_o}{dR_o}$ is same for any intervals $(x_n, x_{n+1})$.

The upper-bound inequality for such interval is given by

$$\int_{x_n}^{x_{n+1}} dR_o\,\{q\varphi_o\} \le \int_{x_n}^{x_{n+1}} dR_o\left\{\frac{1}{w}q^2\right\} \tag{S44}$$

Rearranging Eq.(S44), we get

$$\int_{x_n}^{x_{n+1}} dR_o\,\{R_o^{-l+1}w\varphi_o\} - \int_{x_n}^{x_{n+1}} dR_o\,\{R_o^{-l+1}q\} \le 0 \tag{S45}$$

Multiplying Eq.(S17) by ${R_o}^{-l+1}$ and integrating from $x_n$ to $x_{n+1}$, we get

$$-\int_{x_n}^{x_{n+1}} dR_o\left\{{R_o}^{-l+1}\frac{d}{dR_o}p\frac{d\varphi_o}{dR_o}\right\} + \int_{x_n}^{x_{n+1}} dR_o\left\{{R_o}^{-l+1}w\varphi_o\right\} = \int_{x_n}^{x_{n+1}} dR_o\left\{{R_o}^{-l+1}q\right\} \tag{S46}$$

From the inequality Eq.(S45), we get

$$\int_{x_n}^{x_{n+1}} dR_o\left\{(l-1){R_o}^{-l}p\frac{d\varphi_o}{dR_o}\right\} \le 0 \tag{S47}$$

Since $\frac{d\varphi_o}{dR_o}$ has same sign for all values from $x_n$ to $x_{n+1}$ and $(l-1){R_o}^{-l}p$ are nonnegative for $l = 1,2,\ldots$, $\frac{d\varphi_o}{dR_o} \le 0$. Since this is true for any $x_n$ to $x_{n+1}$, $\frac{d\varphi_o}{dR_o} \le 0$ for any $R_o$.

**Comments on the variational principle**

*Globality of the Minima/Maxima in Eq.(29)*

We provide brief discussion regarding to the justification of the global minima/maxima given by Eq.(29).

More rigorous mathematical proof of the inequality Eq.(29) is beyond the scope of this work, and author refers to the textbook by Arthurs[5] for more detail.

We first note that if the solution to Eq.(8) with the boundary conditions Eq.(9) exists then it is unique.[6] That is, the only admissible function $\varphi$ in the domain of $I[\varphi]$ which makes the first variation of $I[\varphi]$ zero is only $\varphi_o$. Based on this observation, it is clear that $I[\varphi_o]$ is the global maximum provided that the second variation of $I[\varphi]$ is nonpositive (concave down). For if there exist another admissible functions $\varphi'$ that makes $I[\varphi_o] < I[\varphi']$ , then there must exist another stationary point with concave up as $I[\varphi]$ is continuous (for the precise definition of continuity of the functional, see for example Ref.7), which contradict with the uniqueness of the stationary point. Similarly, $G[u_o]$ is the global minimum among all the functions $u$ for the functional $G[u]$ given by Eq.(22) provided that the first variation is zero and second variation is nonnegative.

### Figures

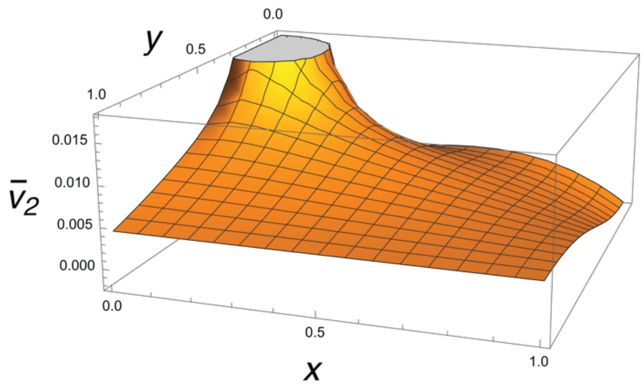


**Figure S1. Plot showing that $v_2(R_o)$ is nonnegative in Eq.(S37).**
For any $R_\text{C}$ , $R_\text{B}$, and $R_o$ in the range $R_\text{C} \le R_o \le R_\text{B}$. $\bar{v}_2(x, y) = (D_\text{trans} R_\text{B}^{-2})^{-1} v_2$ is a unitless function that only involves $x$ and $y$. The condition $R_\text{C} \le R_o \le R_\text{B}$ is equivalent to the condition $0 \le x \le 1$. Furthermore, $y = R_\text{C}/R_\text{B}$ also ranges $0 \le y \le 1$. Therefore, $\bar{v}_l(x, y) \ge 0$ for square unit intervals implies that $v_l(R_o) \ge 0$ .

New York : Oxford University Press, 1980.
(6) Boyce, W. E.; DiPrima, R. C.; Meade, D. B. *Elementary differential equations and boundary value problems*; John Wiley & Sons, 2021.
(7) Gelfand, I. M.; Silverman, R. A. *Calculus of variations*; Courier Corporation, 2000.
